# Motion opponency examined throughout visual cortex with multivariate pattern analysis of fMRI data

**DOI:** 10.1101/2020.05.28.122036

**Authors:** Andrew E. Silva, Benjamin Thompson, Zili Liu

## Abstract

This study explores how the human brain solves the challenge of flicker noise in motion processing. Despite providing no useful directional motion information, flicker is common in the visual environment and exhibits omnidirectional motion energy which is processed by low-level motion detectors. Models of motion processing propose a mechanism called motion opponency that reduces the processing of flicker noise. Motion opponency involves the pooling of local motion signals to calculate an overall motion direction. A neural correlate of motion opponency has been observed in human area MT+/V5 using fMRI, whereby stimuli with perfectly balanced motion energy constructed from dots moving in counter-phase elicit a weaker BOLD response than non-balanced (in-phase) motion stimuli. Building on this previous work, we used multivariate pattern analysis to examine whether the patterns of brain activation elicited by motion opponent stimuli resemble the activation elicited by flicker noise across the human visual cortex. Robust multivariate signatures of opponency were observed in V5 and in V3A. Our results support the notion that V5 is centrally involved in motion opponency and in the reduction of flicker noise during visual processing. Furthermore, these results demonstrate the utility of powerful multivariate analysis methods in revealing the role of additional visual areas, such as V3A, in opponency and in motion processing more generally.

**Highlights:**

- Opponency is demonstrated in multivariate and univariate analysis of V5 data.
- Multivariate fMRI also implicates V3A in motion opponency.
- Multivariate analyses are useful for examining opponency throughout visual cortex.

## Introduction

Motion processing is an essential aspect of vision. However, the successful interpretation of directional motion information is complicated by the presence of flicker noise. Any abrupt change in the luminance of a visual scene, like a flickering light or a bright object appearing suddenly against a dark background, creates flicker noise: omnidirectional and uninformative signals which can be processed just as any true motion signal (Born and Bradley, 2005; Bradley and Goyal, 2008). Therefore, a mechanism to reduce the influence of flicker noise is essential in effective motion processing (Qian et al., 1994).

Classic theoretical models of motion processing employ a mechanism called motion opponency to attenuate the processing of flicker. During motion opponency, a local motion output is calculated by combining all motion signals within the given local area (Adelson and Bergen, 1985; Qian et al., 1994; Reichardt, 1961; Simoncelli and Heeger, 1998; van Santen and Sperling, 1985). The omnidirectional motion signals which define flicker noise are locally balanced and therefore cancel during motion opponency. In contrast, useful motion information is typically directional and not locally balanced. As a result, motion opponency acts as a filter during motion processing, attenuating flicker information while allowing true motion signals to continue for further processing.

Physiological research has identified neural responses indicative of opponency in monkeys. Qian and Andersen (1994) designed a bidirectional and locally motion-balanced dot stimulus in which each randomly-positioned dot was located near a second dot traveling in the opposite direction. This stimulus is now referred to as “counter-phase” (CP) dot motion (Lu et al., 2004). Relative to a bidirectional stimulus without local motion balancing, Qian and Andersen (1994) found that MT neurons exhibited a muted response to counter-phase stimuli. In fact, this response was not significantly greater than the MT response to flicker noise.

Neuroimaging has provided evidence for opponency in human motion processing. Reduced univariate V5 BOLD responses to counter-phase stimuli have been reported in multiple studies and are generally consistent with Qian and Andersen’s (1994) original physiological work (Heeger et al., 1999; Muckli et al., 2002; Thompson et al., 2013). However, suggestions exist that motion opponency and local directional pooling may be distributed throughout the visual cortex in humans (Garcia and Grossman, 2009). Consistent with a multi-region network of local motion pooling, Huck and Heeger (2002) found that relatively high pattern motion-selective responses, indicative of local motion integration, were not exclusive to V5, occurring also in areas V2 and above.

Various non-opponent stimuli have been employed as a comparison against the counterphase stimulus. Often, a stimulus containing the same bidirectional local signals, but without local balancing is employed. One such example of a bidirectional and non-opponent stimulus has been referred to as “in-phase” (IP) (Lu et al., 2004; Silva and Liu, 2018, 2015; Thompson et al., 2013). The in-phase (IP) stimulus is nearly identical to a counter-phase (CP) stimulus, except that both dots within a pair travel in the same direction.

While previous research may be consistent with opponency in the human brain, the human brain’s responses to counter-phase and flicker stimuli have never been directly compared. Because the theoretical formulation of motion opponency selectively reduces flicker noise processing, a more complete understanding of motion opponency in the human brain may be achieved by examining the suppressed response to flicker noise (Adelson and Bergen, 1985; Qian et al., 1994; Reichardt, 1961; Simoncelli and Heeger, 1998; van Santen and Sperling, 1985). If the reduced BOLD response to motion opponent stimuli reported in previous human studies is indeed analogous to theoretical motion opponency, then the human brain may process counter-phase and flicker stimuli similarly. Because both flicker and counter-phase motion stimuli exhibit locally balanced motion, a motion opponent system should output zero net motion in both cases.

With the emergence of multivariate pattern analysis (MVPA) as a powerful tool for understanding neural processing using fMRI (Mahmoudi et al., 2012; Norman et al., 2006; Tong and Pratte, 2012), a detailed exploration of flicker and counter-phase motion processing is now possible. The traditional fMRI region-of-interest analysis involves averaging the responses of all voxels with a region to calculate a single averaged univariate BOLD response. In MVPA classification, a region-wide and voxel-level pattern of activation is inputted, and a classification algorithm predicts which stimulus likely elicited the given brain response. This allows a comparison between stimuli that elicit the same univariate response despite potentially eliciting different patterns of voxel activations.

The current study is the first to examine flicker processing and motion opponency by applying multivariate analysis techniques to fMRI data. Our primary analysis focused on V5. We trained multivariate classifiers with BOLD data associated with in-phase (IP) stimuli, counterphase (CP) stimuli, or non-motion (NM) stimuli exhibiting incoherent onset and offset flicker but no smooth translational movement. Classifiers were trained to discriminate two of the three different stimuli and tested on both trained and untrained stimuli. We predicted the following pattern of results:

1. The multivariate classifier will correctly discriminate IP stimuli from CP and NM stimuli. This result would be consistent with motion opponent processing.
2. The multivariate classifier will systematically misclassify CP stimuli as NM. This result would be consistent with CP stimuli eliciting a similar neural representation to NM due to motion opponency.
3. The multivariate classifier will systematically misclassify NM stimuli as CP. This result would also be consistent with CP stimuli eliciting a similar neural representation to NM due to motion opponency.

As a secondary analysis, we explored the performance of classifiers trained using BOLD data from V1, V2, V3, V3A, and V4 to assess whether BOLD responses indicative of opponency were present throughout the human visual system.

## Material and Methods

### 2.1 Apparatus, Stimuli, and Experimental Procedure

All experimental stimuli were programmed in Python using the Psychopy library (Peirce, 2009, 2007). Stimuli were back-projected onto a screen (12 cm × 9 cm useable area, 1024 × 768 resolution, 60 hz refresh rate) that was mounted above the fMRI head coil. Participants viewed the display through a mirror. Due to differences in head size, viewing distances ranged between 22 cm and 25 cm, and therefore the size of one pixel ranged between 0.027 and 0.031 degrees. All stimuli were presented in front of a solid gray background (luminance 4 cd/m^2^).

Participants fixated on a central black square dot 5 pixels in size throughout an entire experimental run. The visual presentation alternated between a 12-second stimulus block and a 12-second blank block displaying only the fixation point. During the stimulus blocks, 250 pairs of randomly distributed white square dots (luminance 67 cd/m^2^) of size 3 pixels were presented. Each dot was initially placed no more than 8 pixels away from its paired partner along a common orientation, creating a Glass pattern (Glass, 1969). The Glass pattern could be oriented either horizontally or vertically in any given block. All dots had a limited dot lifetime of 150 ms before being randomly replotted.

To engage attention, a mildly effortful behavioral task was employed. Stimulus blocks were divided into 6 trials, each lasting 1.1 seconds. On each trial, the Glass pattern orientation was 15 degrees clockwise or counterclockwise from the block’s overall cardinal orientation. Each block contained 3 clockwise and 3 counterclockwise trials. Participants indicated the orientation of each trial using a button response box. All participants achieved ceiling performance. An inter-trial interval of 500 ms was used, during which no dots were presented.

Three different paired-dot stimulus conditions were presented separately in blocks that were randomly interleaved throughout each scanning run. Each block could be composed of counter-phase (CP), non-motion (NM), or in-phase (IP) stimuli. During counter-phase blocks, the two dots in a pair travelled in opposite directions. CP pairs were initially separated by 8 pixels along the Glass pattern orientation and traveled toward one another, crossed, and were randomly replotted after again achieving a separation of 8 pixels. To temporally stagger the replotting of CP dots, each CP pair was initially plotted at a randomly selected point along its full trajectory.

During IP blocks, both dots within a pair traveled in the same direction along the orientation of the Glass pattern. Each pair was independently assigned a random initial lifetime to temporally stagger the replotting of dot pairs, and each pair was independently assigned a random within-pair distance between 0 and 8 pixels. Different IP pairs travelled in opposite directions along the Glass pattern orientation, creating a bidirectional stimulus. Because IP and CP dots all traveled 8 pixels during their 150ms limited lifetime, the dot speed ranged from 2.3 to 2.6 degrees/s, depending on viewing distance.

NM dots behaved identically to in-phase dots, except that there was no translational motion. Critically, in-phase and counter-phase stimuli contained the same number of left and right motion signals, and the Glass patterns of all three conditions were indistinguishable from one another. One experimental run contained 6 blocks of each paired-dot condition, totaling 18 blocks per run. Each participant performed 8 runs, totaling 144 blocks (48 blocks per condition). Stimulus diagrams are presented in Figure 1.

**Figure 1.**
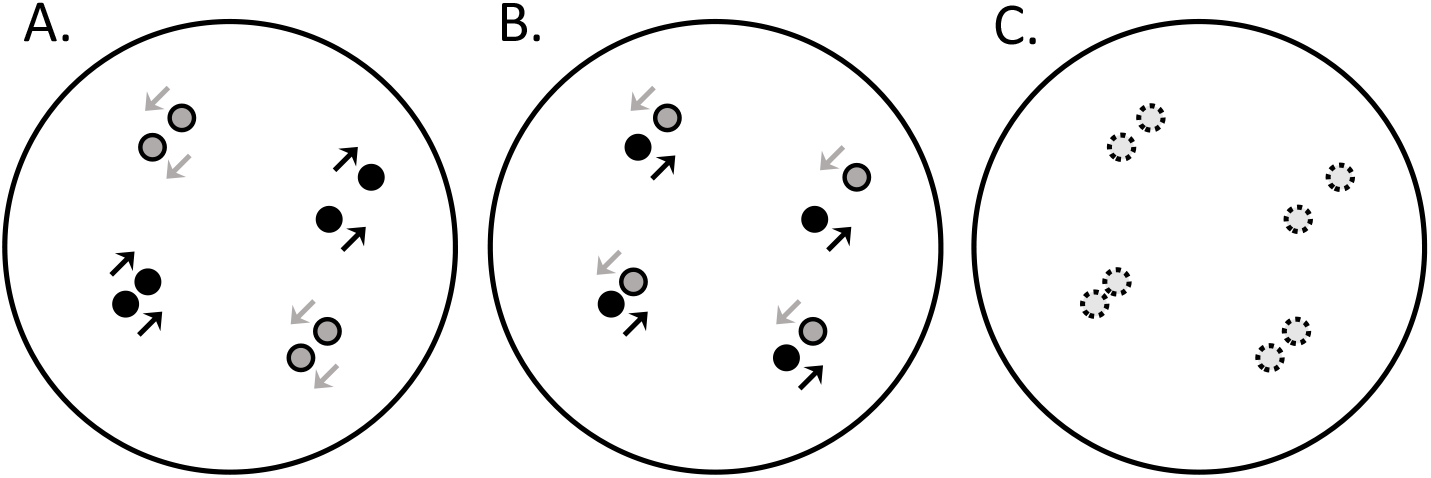
Diagrams of in-phase (A), counter-phase (B), and non-motion stimuli (C). Dots are shaded according to their direction of motion. Non-motion dots are represented by broken circles. All dots had limited lifetimes and the stimuli exhibited indistinguishable Glass patterns.

### 1.2 Participants

Functional neuroimaging data were collected from five participants. All participants had normal or corrected-to-normal vision. Informed consent was obtained, and all participants were treated in accordance with the Code of Ethics of the World Medical Association (Declaration of Helsinki). For their participant, participants received CAN$200 (CAN$50 per scanning hour).

### 2.3 Magnetic Resonance Imaging

All scans took place in the Centre for Functional and Metabolic Mapping at the University of Western Ontario’s Robarts Research Institute on the 7T Siemens Magnetom scanner. All functional scans used an 8-channel transmit, 32-channel receive coil optimized for the occipital pole and providing an unobstructed field-of-view to the visual stimulus. All anatomical scans used an 8-channel transmit, 32-channel whole-head coil. For each participant, we collected an anatomical scan (MP2RAGE, 224 sagittal slices, 0.7 mm isotropic voxel size, TR=6000 ms, TE= 2.73 ms, Flip angle 1 = 4°, Flip angle 2 = 5°, TI 1 =800 ms, TI 2 = 2700 ms), two retinotopy scans, one with rotating wedge stimuli and one with expanding ring stimuli (Sixty coronal slices originating at the posterior pole, 1.5 mm isotropic voxel size, TR = 1000 ms, TE = 19.6 ms, Flip angle = 45°, slice order: interleaved, phase condition direction: FH, pulse: gradient echo, imaging type: EPI), one V5 localizer scan (Sixty coronal slices originating at the posterior pole, 1.5 mm isotropic voxel size, TR = 1600 ms, TE = 19.6 ms, Flip angle = 45°, slice order: interleaved, phase condition direction: FH, pulse: gradient echo, imaging type: EPI), and eight experimental scans (Sixty coronal slices originating at the posterior pole, 1.5 mm isotropic voxel size, TR = 1200 ms, TE = 19.6 ms, Flip angle = 45°, slice order: interleaved, phase condition direction: FH, pulse: gradient echo, imaging type: EPI).

### 2.4 Preprocessing

fMRI data were preprocessed in BrainVoyager. The functional localizer analysis and the univariate experimental analysis were conducted using BrainVoyager QX 2.8.4 (Formisano et al., 2006; Goebel et al., 2006). Functional data were preprocessed using motion correction, slice scan time correction, and highpass filtering. The functional scans were coregistered to the native anatomical space using BrainVoyager’s coregistration and visualization tools. All transformations were applied with sinc interpolation.

### 2.5 Regions of Interest Localization

A standard rotating wedge and expanding ring retinotopic mapping procedure was used to identify areas V1, V2, V3, V3A, and V4 (Engel et al., 1997; Sereno et al., 1995). The black-and-white checkerboard wedges spanned 45°, shifted 11.25° per TR (1000 ms) and completed 7 full cycles during the session. The checkerboard rings began centrally and expanded into the periphery once per TR (1000 ms). Twenty such expansions per cycle occurred, and 7 full cycles were completed during the session. The largest ring had an outer radius of 384 pixels (between 10.4° and 11.9°) and an inner radius of 270 pixels (between 7.3° and 8.4°). The retinotopic stimuli flickered and reversed their contrast polarity at a rate of 8 hz. V5 localization stimuli were composed of 1,348 white square dots with a side length of 3 pixels alternating between inward and outward radial motion. The dots traveled 4 pixels per frame (between 6.5°/s and 7.4 °/s) and reversed direction every 2 seconds. Four 16-second blocks were presented, alternating with 16-second blank periods containing completely static dots exhibiting no limited lifetime. In every localization scan, participants performed a fixation task, indicating when the central fixation randomly alternated between “O” and “X”.

Bilateral V5 was identified for each participant. First, a GLM was fit to the V5 localization data using a box-car stimulus model and BrainVoyager’s default double-gamma HRF. The model additionally contained *z*-scored head-motion nuisance regressors. A wholebrain, voxel-wise contrast of moving dots verses static dots was applied (FDR, *q* < 0.05). V5 was defined as significant clusters of voxels bilaterally located near the ascending limb, or the posterior continuation, of the inferior temporal sulcus or the posterior bank of the superior temporal sulcus (Dumoulin et al., 2000).

To identify areas V1-V4, a 3D brain surface model was constructed from the skullstripped and Talairach-transformed anatomical scan in BrainVoyager. The surface was inflated, cut across the calcarine sulcus, flattened, and corrected for surface distortions. A whole-brain, voxel-wise cross-correlation analysis was carried out and mapped onto the flattened brain surface, and the borders of V1-V4 were identified by observing the cross-correlation polarity reversals running along the calcarine sulcus. A GLM was then fit to all experimental data collected for the participant, and a voxel-wise contrast of stimulus period verses blank period was applied (FDR, *q* < 0.05). The final ROIs were defined as the significant voxels within the ROI borders.

### 2.6 fMRI analysis

Model-independent univariate BOLD time courses were extracted for each paired-dot condition for each visual area from the Talairach-space transformed data. The data were then transformed to native functional space for the MPVA analysis. Within-subject multivariate pattern classification analyses were carried out on fitted voxel-wise GLM betas for each visual area using support vector machines. Betas were calculated using the MATLAB toolbox NeuroElf (www.neuroelf.net). One GLM per ROI was fitted with each individual block as a separate regressor (Mumford et al., 2012; Rissman et al., 2004) and with *z*-scored head-motion data as additional nuisance regressors. Each block was modeled as an individual box-car, convolved with NeuroElf’s default double-gamma HRF and *z*-score normalized. Blocks containing different cardinal orientations (vertical vs. horizontal) were collapsed and analyzed together.

The classification analysis utilized Linear Support Vector Machines programmed in Python with the Scikit-learn library using the default hyperparameter settings (Pedregosa et al., 2011). Three independent classifiers were trained to discriminate between, and then tested on, IP and CP, IP and NM, and CP and NM blocks. All SVMs utilized 8-fold cross validation, whereby the classifier was trained on 7 of 8 runs and tested on the remaining run. This occurred 8 times per SVM such that each run, in turn, served as the testing set, and the final performance was the average of all 8 folds. Data from all identified ROIs were analyzed in this way.

A further classification analysis was carried out on data from visual area ROIs exhibiting greater than 70% group-mean accuracy in at least two of the three pairwise classifiers. Three pairwise SVMs were trained identically as in the previous analysis, but the testing dataset was composed of data from the untrained condition. For example, the classifier trained on 7 folds of IP and CP data was tested on the NM data of the 8^th^ fold, and an 8-fold cross validation scheme was again used. This analysis was used to probe for the presence of any systematic misclassification bias between in-phase, counter-phase, and non-motion conditions. Because this analysis is unlikely to uncover systematic classification bias if the previous condition discrimination analysis performs poorly, it was only carried out with data from ROIs exceeding 70% classification accuracy in two condition discrimination classifiers.

In all multivariate analyses, significance was established using within-subject permutation tests in which the condition labels were randomly permuted within runs, ensuring that each run preserved the same number of each type of label (Etzel and Braver, 2013). A total of 15,200 permuted datasets were tested. All participants received the same permutations, and the results of each permutation were averaged across participants to create one group null distribution against which the group average performance could be compared (Etzel, 2015). The permuted *ρ* was defined as the percentile rank of the non-permuted group average divided by the total number of permutations + 1 (15,201).

## Results

### 3.1 Univariate Results

Figure 2 shows the average BOLD time-series of IP, CP, and NM blocks, collapsed across participants and percent-normalized by the voxel intensity of the first TR at stimulus onset. Consistent with previous studies, visual inspection reveals an increased IP BOLD response at area V5 and mostly overlapping activity across all three conditions for the other visual areas tested.

**Figure 2.**
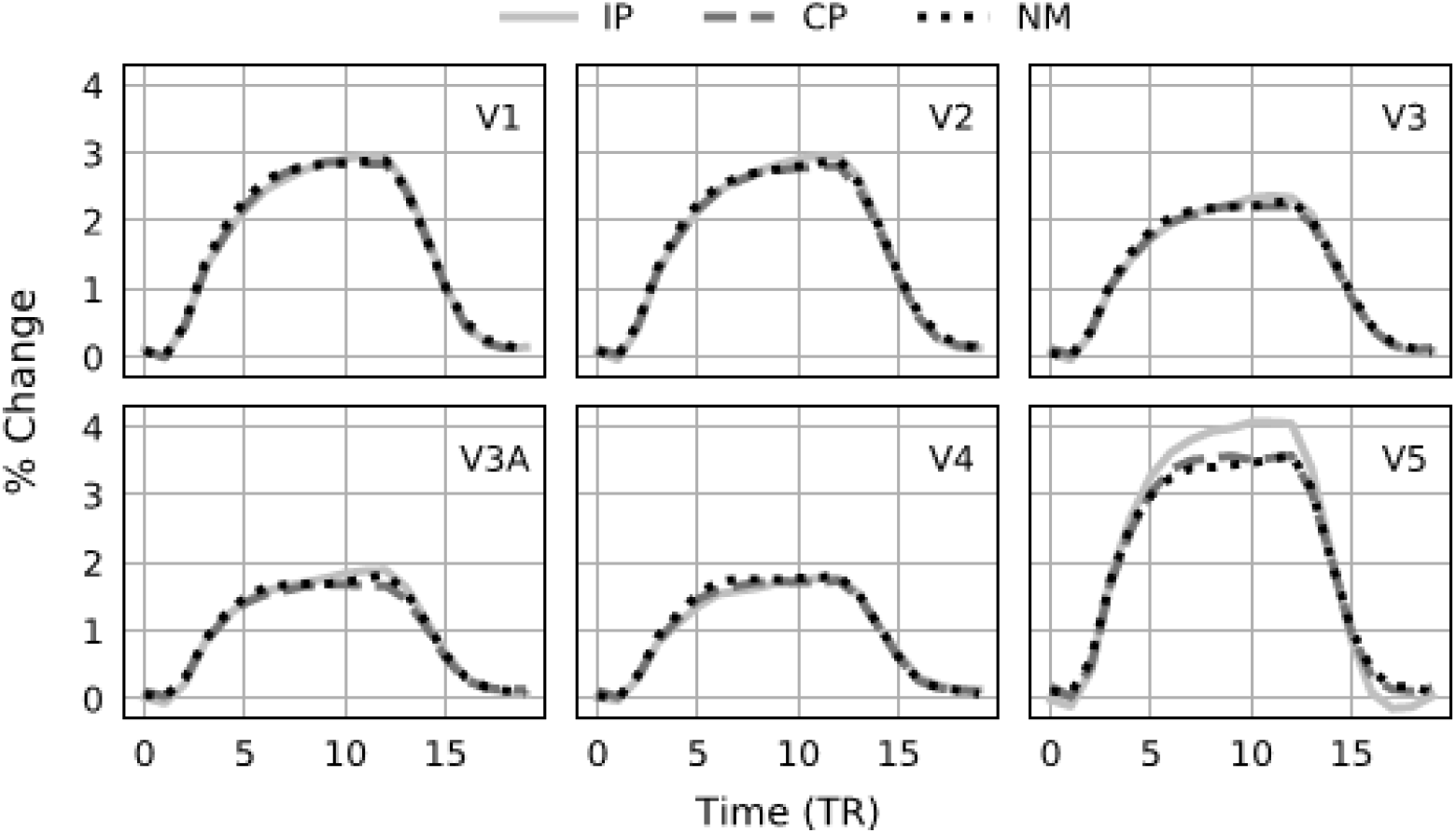
Group-averaged univariate time-series during IP, counter-phase CP, and NM blocks plotted as percent change for all ROIs. Data were normalized relative to the onset of the visual stimulus (TR = 0).

### 3.2 Multivariate Results

For all MVPA analyses, significance was determined using a permutation test with 15,200 random permutations. Therefore, the minimum *p* possible is 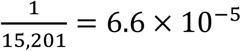 when the true value is more extreme than every null value.

IP, CP, and NM blocks were used to train and test IP v. CP, IP v. NM, and CP v. NM classifiers to examine the separability of each condition with a one-tailed permutation test. A Bonferroni correction for multiple comparisons was applied. This analysis contains 3 comparisons across 6 ROIs; therefore, a critical *p* of 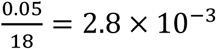 was set to determine better-than-chance accuracy.

Every ROI achieved greater than chance performance when discriminating IP and CP. In increasing performance order: V1 – 57%, *p* = 9.2 × 10^-4^; V4 – 58%, *p* = 3.3 × 10^-4^; V2 – 62%, *p* = 6.6 × 10^-5^; V3 – 65%, *p* = 6.6 × 10^-5^; V3A – 76%, *p* = 6.6 × 10^-5^; V5 – 79%, *p* = 6.6 × 10^-5^. Area V1 failed to achieve the significance cutoff when discriminating IP and NM: 56%, *p* = 4.4 × 10^-3^. However, all other areas successfully discriminated IP and NM. In increasing performance order: V2 – 60%, *p* = 1.3 × 10^-4^; V4 – 61%, *p* = 6.6 × 10^-5^; V3 – 67%, *p* = 6.6 × 10^-5^; V5 – 75%, *p* = 6.6 × 10^-5^; V3A – 81%, *p* = 6.6 × 10^-5^. When discriminating CP and NM, performance was relatively poorer across all ROIs. Only data from areas V3 and V3A surpassed the threshold of significance, V3 – 58%, *p* = 6.6 × 10^-5^; V3A – 59%, *p* = 6.6 × 10^-5^. The remaining ROIs did not achieve significance when discriminating CP and NM, V5 – 52%, *p* = 1.5 × 10^-1^; V1 – 54%, *p* = 3.0 × 10^-2^; V2 – 54%, *p* = 3.5 × 10^-2^; V4 – 55%, *p* = 4.5 × 10^-3^. The condition discrimination results for all ROIs are plotted in Figure 3.

**Figure 3.**
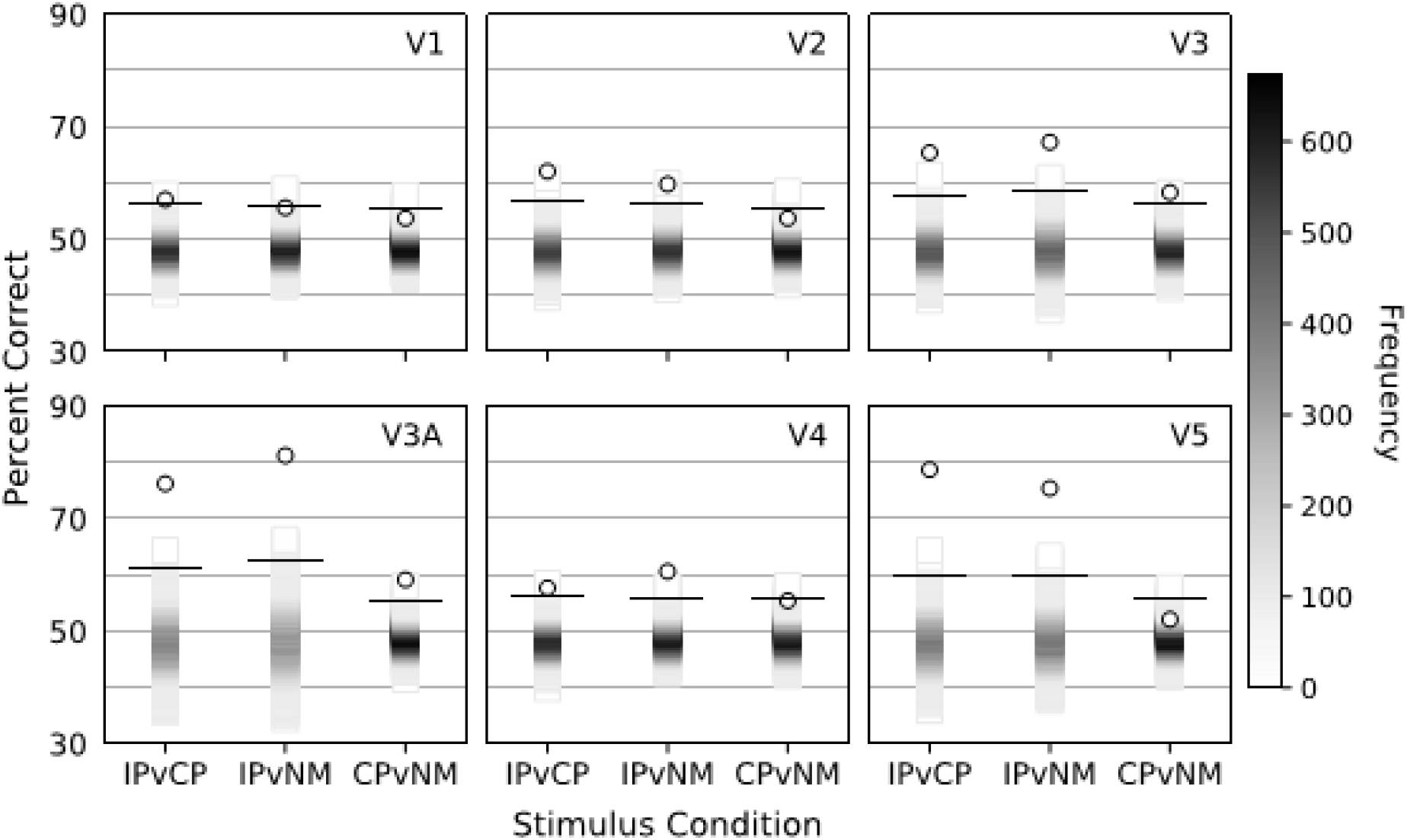
Condition discrimination MVPA results plotted in percent correct. IPvCP plots the discrimination between in-phase and counter-phase. IPvNM plots the discrimination between in-phase and non-motion. CPvNM plots the discrimination between counter-phase and non-motion. The shaded bar represents the estimated null distribution. Darker shades represent a higher frequency of values achieving the associated percent correct. The null distribution was estimated using 15,200 permutations. Performance using the unpermuted dataset is plotted as circles. The lines illustrate the performance required to exceed the critical *p* of 2.8 × 10^-3^.

Because the data from V3A and V5 achieved at least 70% accuracy in two comparisons, their data were submitted to a further analysis designed to probe for any systematic misclassification biases. Three classifiers were trained to discriminate between IP and CP, IP and NM, and CP and NM identically as in the previous analysis, but they were tested with the untrained condition to determine whether the untrained condition would elicit a consistent misclassification error. A two-tailed permutation test was used to determine significance, and a Bonferoni correction for multiple comparisons was applied. This analysis contains 3 comparisons across 2 ROIs; therefore, a critical *p* of 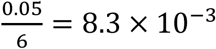 was set to determine better-than-chance performance.

The classier trained to discriminate CP and NM was tested with IP data and there was no systematic misclassification of IP in either V3A or V5 (See Figure 4A). However, a significant misclassification bias was found with the classifier trained to discriminate IP and NM and tested with CP data. CP data was classified more often as NM in both ROIs; V3A – 73% classified as NM, *p* = 6.6 × 10^-5^; and V5 – 72% classified as NM, *p* = 6.6 × 10^-5^ (See figure 4B). Similarly, a significant misclassification bias was found with the classifier trained to discriminate IP and CP and tested with NM data. NM data was classified more often as CP in both ROIs; V3A – 71% classified as CP, *p* = 1.3 × 10^-4^; and V5 – 68% classified as CP, *p* = 3.9 × 10^-4^ (See Figure 4C). See Supp. Figure A for a plot of misclassification bias across all ROIs.

**Figure 4:**
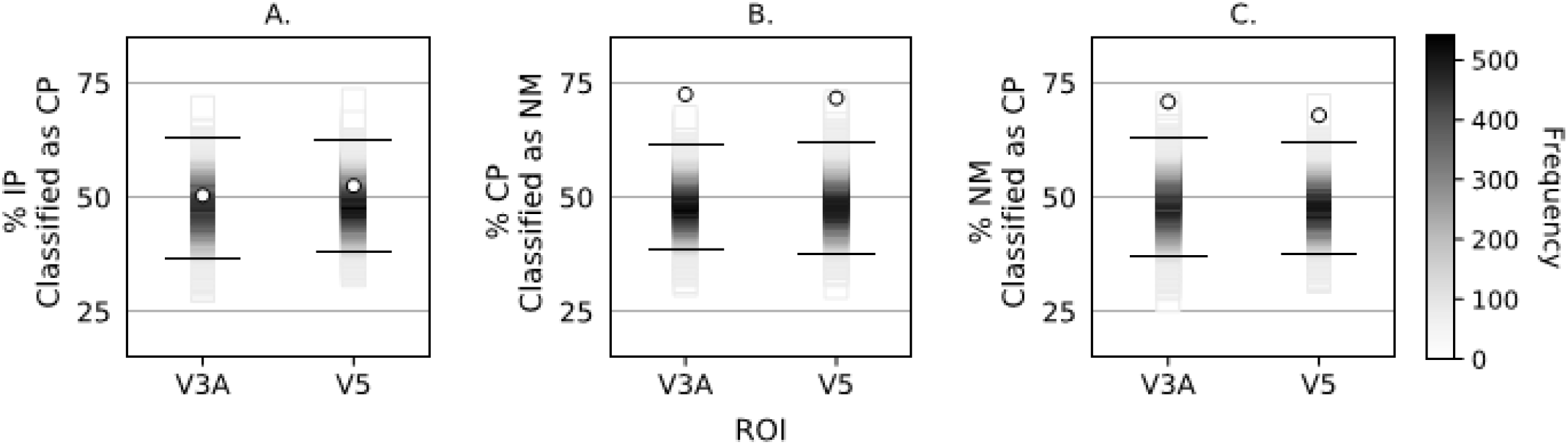
Misclassification bias analysis. A. The results of the classifier trained to discriminate CP and NM and tested with NM data, plotted as the percent of NM blocks classified as CP. B. The results of the classifier trained to discriminate IP and NM and tested with CP data, plotted as the percent of CP blocks classified as NM. C. The results of the classifier trained to discriminate IP and CP and tested with NM data, plotted as the percent of NM blocks classified as CP. Plotting conventions are identical to Figure 2. Because this analysis involves 6 comparisons, the lines illustrate the performance required to exceed the critical *p* of 8.3 × 10^-3^.

## General Discussion

The current study examined the human motion opponency system using a novel non-motion flicker-based stimulus and a multivariate analysis of fMRI data. Motion opponency involves the pooling of local motion signals to output an overall motion direction and is therefore useful in flicker noise reduction (Adelson and Bergen, 1985; Qian et al., 1994; Reichardt, 1961; Simoncelli and Heeger, 1998; van Santen and Sperling, 1985). As a result, a motion-opponent system may process counter-phase motion and flicker noise similarly. We therefore hypothesized that BOLD data from any visual area involved in opponency would elicit a specific multivariate signature: 1. strong separability of in-phase data, and 2. systematic misclassification of counterphase blocks as non-motion and non-motion blocks as counter-phase.

Previous neuroimaging work reported suppressed univariate counter-phase V5 responses (Garcia and Grossman, 2009; Heeger et al., 1999; Muckli et al., 2002; Thompson et al., 2013). The current study directly extended this result by comparing the counter-phase and non-motion flicker responses to each other as well as to the in-phase response. Our multivariate predictions were fully born out within V3A and V5. These results are consistent with the notion that V5 similarly processes counter-phase and flicker stimuli and that motion opponent suppression is recruited during the processing of both stimuli.

Motion opponency is typically associated with area V5/MT receiving inputs from V1 (Bradley and Goyal, 2008; Qian and Andersen, 1994). However, the present results also suggest involvement of V3A in motion opponency, finding robust multivariate signals of opponency in V3A. Because opponency contributes a necessary noise-reduction step in motion processing, the suggestion that V3A participates in opponency may be consistent with previous findings that V3A participates in motion processing and cooperates with V5 in motion perceptual learning (Braddick et al., 2001; Chen et al., 2016, 2015; Tootell et al., 1997).

It might also be noted that unlike area V5, the V3A univariate in-phase, counter-phase, and non-motion timeseries curves appear to overlap (See Figure 2). The visually distinct appearance of the univariate V5 time series may suggest a qualitatively different or more central role in opponency. Another possibility is that the population of V3A neurons participating in opponency is too small to be visually apparent in the univariate BOLD response. Multivariate classification methods are more powerful than univariate methods (Mur et al., 2009; Tong and Pratte, 2012), and the presence of motion opponency throughout the visual system might be more reliably detected with these more powerful methods.

The results of the current study strengthen the idea that both counter-phase and flicker stimuli elicit motion opponency in the human brain. Furthermore, they demonstrate that multivariate analyses are powerful tools to examine motion opponency throughout the visual system, providing evidence that area V3A may participate in motion opponency alongside V5.

Further work is required to clarify V3A’s potential role in motion opponency and to examine the full motion opponent network in the human brain.

## Funding

This work is supported by NSF Graduate Research Fellowship Grant Research No. DGE-11444087 to A.E.S., NSERC Grants RPIN-05394 and RGPAS-477166 to B.T, and NSF Grant No. 0617628 to Z.L.

**Figure A:**
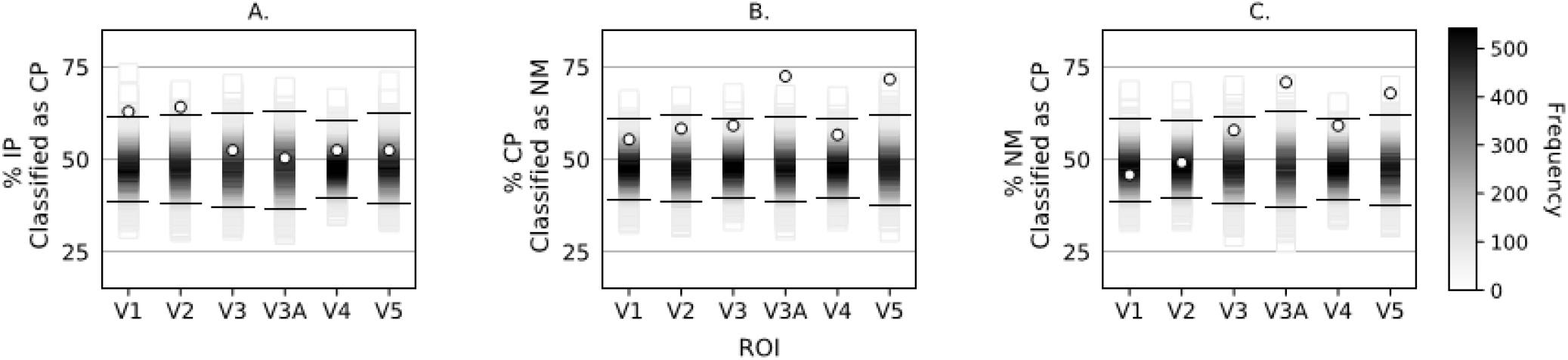
Misclassification bias analysis for every ROI, plotted as in Figure 3. The line illustrates the performance required to exceed *p* = 8.3 × 10^-3^, the critical *p* used in the text. If a Bonferroni adjustment for 18 comparisons were applied instead, the critical *p* would be 2.8 × 10^-3^ and the only significant values would be V3A and V5 in plots B. and C.

## References

Adelson, E.H., Bergen, J.R., 1985. Spatiotemporal energy models for the perception of motion. J. Opt. Soc. Am. A 2, 284–299. https://doi.org/10.1364/JOSAA.2.000284

Born, R.T., Bradley, D.C., 2005. Structure and function of visual area MT. Annu. Rev. Neurosci. 28, 157–189. https://doi.org/10.1146/annurev.neuro.26.041002.131052

Braddick, O.J., O’Brien, J.M.D., Wattam-Bell, J., Atkinson, J., Hartley, T., Turner, R., 2001. Brain Areas Sensitive to Coherent Visual Motion. Perception 30, 61–72. https://doi.org/10.1068/p3048

Bradley, D.C., Goyal, M.S., 2008. Velocity computation in the primate visual system. Nat. Rev. Neurosci. 9, 686–695. https://doi.org/10.1038/nrn2472

Chen, N., Bi, T., Zhou, T., Li, S., Liu, Z., Fang, F., 2015. Sharpened cortical tuning and enhanced cortico-cortical communication contribute to the long-term neural mechanisms of visual motion perceptual learning. Neuroimage 115, 17–29. https://doi.org/10.1016/j.neuroimage.2015.04.041

Chen, N., Cai, P., Zhou, T., Thompson, B., Fang, F., 2016. Perceptual learning modifies the functional specializations of visual cortical areas. Proc. Natl. Acad. Sci. 113, 5724–5729. https://doi.org/10.1073/pnas.1524160113

Dumoulin, S.O., Bittar, R.G., Kabani, N.J., Baker, C.L., Le Goualher, G., Pike, G.B., Evans, A.C., 2000. A New Anatomical Landmark for Reliable Identification of Human Area V5/MT: a Quantitative Analysis of Sulcal Patterning. Cereb. Cortex 10, 454–463. https://doi.org/10.1093/cercor/10.5.454

Engel, S.A., Glover, G.H., Wandell, B.A., 1997. Retinotopic organization in human visual cortex and the spatial precision of functional MRI. Cereb. Cortex 7, 181–192. https://doi.org/10.1093/cercor/7.2.181

Etzel, J.A., 2015. MVPA Permutation Schemes: Permutation Testing for the Group Level, in: 2015 International Workshop on Pattern Recognition in Neuroimaging. IEEE, pp. 65–68. https://doi.org/10.1109/PRNI.2015.29

Etzel, J.A., Braver, T.S., 2013. MVPA Permutation Schemes: Permutation Testing in the Land of Cross-Validation, in: 2013 International Workshop on Pattern Recognition in Neuroimaging. IEEE, pp. 140–143. https://doi.org/10.1109/PRNI.2013.44

Formisano, E., Di Salle, F., Goebel, R.W., 2006. Fundamentals of data analysis methods in fMRI, in: Landini, L., Positano, V., Santarelli, M.F. (Eds.), Advanced Image Processing in Magenetic Resonance Imaging. Marcel Dekker, New York, pp. 481–504.

Garcia, J.O., Grossman, E.D., 2009. Motion opponency and transparency in the human middle temporal area. Eur. J. Neurosci. 30, 1172–1182. https://doi.org/10.1111/j.1460-9568.2009.06893.x

Garcia, J.O., Grossman, E.D., 2008. Necessary but not sufficient: Motion perception is required for perceiving biological motion. Vision Res. 48, 1144–1149. https://doi.org/10.1016/j.visres.2008.01.027

Glass, L., 1969. Moiré Effect from Random Dots. Nature 223, 578–580. https://doi.org/10.1038/223578a0

Goebel, R., Esposito, F., Formisano, E., 2006. Analysis of functional image analysis contest (FIAC) data with brainvoyager QX: From single-subject to cortically aligned group general linear model analysis and self-organizing group independent component analysis. Hum. Brain Mapp. 27, 392–401. https://doi.org/10.1002/hbm.20249

Heeger, D.J., Boynton, G.M., Demb, J.B., Seidemann, E., Newsome, W.T., 1999. Motion opponency in visual cortex. J. Neurosci. 19, 7162–7174.

Huk, A.C., Heeger, D.J., 2002. Pattern-motion responses in human visual cortex. Nat. Neurosci. 5, 72–75. https://doi.org/10.1038/nn774

Lu, H., Qian, N., Liu, Z., 2004. Learning motion discrimination with suppressed MT. Vision Res. 44, 1817–1825. https://doi.org/10.1016/j.visres.2004.03.002

Mahmoudi, A., Takerkart, S., Regragui, F., Boussaoud, D., Brovelli, A., 2012. Multivoxel pattern analysis for fMRI data: A review. Comput. Math. Methods Med. 2012. https://doi.org/10.1155/2012/961257

Muckli, L., Singer, W., Zanella, F.E., Goebel, R., 2002. Integration of multiple motion vectors over space: an fMRI study of transparent motion perception. Neuroimage 16, 843–856. https://doi.org/10.1006/nimg.2002.1085

Mumford, J.A., Turner, B.O., Ashby, F.G., Poldrack, R.A., 2012. Deconvolving BOLD activation in event-related designs for multivoxel pattern classification analyses. Neuroimage 59, 2636–43. https://doi.org/10.1016/j.neuroimage.2011.08.076

Mur, M., Bandettini, P.A., Kriegeskorte, N., 2009. Revealing representational content with pattern-information fMRI — an introductory guide. Soc. Cogn. Affect. Neurosci. 4, 101–109. https://doi.org/10.1093/scan/nsn044

Norman, K.A., Polyn, S.M., Detre, G.J., Haxby, J. V., 2006. Beyond mind-reading: multi-voxel pattern analysis of fMRI data. Trends Cogn. Sci. 10, 424–430. https://doi.org/10.1016/j.tics.2006.07.005

Pedregosa, F., Varoquaux, G., Gramfort, A., Michel, V., Thirion, B., Grisel, O., Blondel, M., Müller, A., Nothman, J., Louppe, G., Prettenhofer, P., Weiss, R., Dubourg, V., Vanderplas, J., Passos, A., Cournapeau, D., Brucher, M., Perrot, M., Duchesnay, É., 2011. Scikit-learn: Machine Learning in Python. J. Mach. Learn. Res. 12, 2825–2830.

Peirce, J.W., 2009. Generating Stimuli for Neuroscience Using PsychoPy. Front. Neuroinform. 2, 10. https://doi.org/10.3389/neuro.11.010.2008

Peirce, J.W., 2007. PsychoPy-Psychophysics software in Python. J. Neurosci. Methods 162, 8–13. https://doi.org/10.1016/j.jneumeth.2006.11.017

Qian, N., Andersen, R.A., 1994. Transparent motion perception as detection of unbalanced motion signals. II. Physiology. J. Neurosci. 14, 7367–7380.

Qian, N., Andersen, R.A., Adelson, E.H., 1994. Transparent motion perception as detection of unbalanced motion signals. III. Modeling. J. Neurosci. 14, 7381–7392.

Reichardt, W., 1961. Autocorrelation, a Principle for the Evaluation of Sensory Information by the Central Nervous System, in: Rosenblith, W.A. (Ed.), Sensory Communication. Wiley, New York.

Rissman, J., Gazzaley, A., D’Esposito, M., 2004. Measuring functional connectivity during distinct stages of a cognitive task. Neuroimage 23, 752–763. https://doi.org/10.1016/J.NEUROIMAGE.2004.06.035

Sereno, M., Dale, A., Reppas, J., Kwong, K., Belliveau, J., Brady, T., Rosen, B., Tootell, R., 1995. Borders of multiple visual areas in humans revealed by functional magnetic resonance imaging. Science (80-.). 268, 889–893. https://doi.org/10.1126/science.7754376

Silva, A.E., Liu, Z., 2018. Spatial proximity modulates the strength of motion opponent suppression elicited by locally paired dot displays. Vision Res. 144, 1–8. https://doi.org/10.1016/j.visres.2018.01.004

Silva, A.E., Liu, Z., 2015. Opponent backgrounds reduce discrimination sensitivity to competing motions: Effects of different vertical motions on horizontal motion perception. Vision Res. 113, 55–64. https://doi.org/10.1016/j.visres.2015.05.007

Simoncelli, E.P., Heeger, D.J., 1998. A model of neuronal responses in visual area MT. Vision Res. 38, 743–761. https://doi.org/10.1016/S0042-6989(97)00183-1

Thompson, B., Tjan, B.S., Liu, Z., 2013. Perceptual learning of motion direction discrimination with suppressed and unsuppressed MT in humans: an fMRI study. PLoS One 8, e53458. https://doi.org/10.1371/journal.pone.0053458

Tong, F., Pratte, M.S., 2012. Decoding Patterns of Human Brain Activity. Annu. Rev. Psychol. 63, 483–509. https://doi.org/10.1146/annurev-psych-120710-100412

Tootell, R.B.H., Mendola, J.D., Hadjikhani, N.K., Ledden, P.J., Liu, A.K., Reppas, J.B., Sereno, M.I., Dale, A.M., 1997. Functional Analysis of V3A and Related Areas in Human Visual Cortex. J. Neurosci. 17, 7060–7078. https://doi.org/10.1523/JNEUROSCI.17-18-07060.1997

van Santen, J.P.H., Sperling, G., 1985. Elaborated Reichardt detectors. J. Opt. Soc. Am. A 2, 300–321. https://doi.org/10.1364/JOSAA.2.000300

